# The 16p11.2 microdeletion exacerbates neurodevelopmental alterations induced by early-life microbiome perturbation

**DOI:** 10.1101/2025.02.25.639888

**Authors:** Courtney R. McDermott, Zhan Gao, Anya S. Mirmajlesi, Christiana Ntim, Katherine Kimbark, Divya Thomas, Zain Mughal, Xue-Song Zhang, Xiaofeng Zhou, Daniel Popov, Alisa Halchenko, Jinchuan Xing, Smita Thakker-Varia, Janet Alder, James H. Millonig, Benjamin A. Samuels, Martin J. Blaser, Emanuel DiCicco-Bloom

## Abstract

Neurodevelopmental disorders (NDDs) arise from interactions between genetic factors and environmental exposures, with infancy representing a critical period of vulnerability. This exploratory, preclinical study investigated whether the 16p11.2 microdeletion (16pDel), a NDD-associated genetic variant, exacerbates the effects of early-life therapeutic antibiotic exposure on the gut microbiome, hippocampal development, and behavior. Cefdinir, selected for its epidemiological association with NDD risk, acutely perturbed the gut microbiome, causing sustained reductions in *Lachnospiraceae*. These changes were followed by alterations in sociability, risk assessment, and associative learning. Notably, only cefdinir-exposed 16pDel mice exhibited altered hippocampal stem cell dynamics and gene expression, demonstrating genotype-dependent susceptibility. Increased intestinal permeability and alterations in arginine biosynthesis and glycerophospholipid metabolism implicate gut barrier dysfunction as a contributing factor. Our findings suggest that genetic composition can exacerbate neurodevelopmental consequences of early-life microbiome perturbations, identify metabolic pathways for potential interventions, and support cautious antibiotic use during infancy, especially in genetically vulnerable populations.

## Introduction

The average child in the United States receives more than two courses of antibiotics by age three, which are often prescribed empirically rather than in response to a defined bacterial infection^1,2^. This over- and misuse of antibiotics during a sensitive time in life has been linked to adverse health outcomes, including neurodevelopmental disorders (NDDs)^3–6^. For example, birth cohort studies have shown associations of early-life antibiotic exposure with increased risks of attention deficit hyperactivity disorder, developmental delay, learning disability, and autism spectrum disorder (ASD)^4–8^.

Furthermore, our recent study of 14,572 children found that cephalosporin exposure in the first two years of life was significantly associated with increased ASD risk in both males and females (hazard ratios 1.89 and 2.77, respectively)^3^. Although these studies show association rather than causation, they warrant further study into the underlying factors linking early-life antibiotic use to NDD outcomes.

Accumulating evidence suggests that the gut microbiome may be a nexus between antibiotic exposure and NDDs^9–15^. Studies in both humans and experimental animals have shown that antibiotic exposure disrupts gut microbiome homeostasis, as indicated by altered bacterial richness and community structure^16–19^. Such alterations, particularly in early life, are hypothesized to affect neurodevelopment through the gut microbiome-brain axis, which integrates vagal, metabolic, endocrine, and immune signaling pathways^20–23^. However, most research on neural and behavioral changes following antibiotic exposure has focused primarily on the adult brain, overlooking the sensitive neurodevelopmental timeframe when neural populations and connectivity are established. Such studies have used prolonged, high-dose antibiotic regimens to suppress the microbiome, an approach that does not reflect typical pediatric clinical practice^10,12,24–29^. Similarly, comparisons of human children with and without ASD cannot address issues of causal roles^30^. To address these gaps, we explored how brief, clinically relevant antibiotic exposure during early life in mice influence the gut microbiome and fundamental stages of neurodevelopment within the hippocampus.

In the postnatal rodent brain, the hippocampal dentate gyrus is a critical site of neurogenesis and astrogenesis, where neural stem cell (NSC) proliferation peaks during the first two weeks of life^31^. These NSCs give rise to intermediate progenitor cells that frequently divide before migrating and differentiating into neurons or astrocytes, both essential for proper neural circuitry^32^. NSCs are highly sensitive to both inherited and environmental factors, and alterations in their dynamics have been implicated in NDD pathogenesis^33–36^. Although we hypothesize that early-life antibiotic exposure impacts neurodevelopment, only a subset of exposed infants develop NDDs, suggesting that genetic composition may modulate susceptibility to microbial perturbation. To address this, we compared 16p11.2 microdeletion copy number variation (16pDel CNV) and wildtype (WT) mice. Deletions or duplications at the 16p11.2 locus represent one of the most frequently observed genetic mutations associated with NDDs^37,38^. An ASD diagnosis occurs in ∼30% of children with 16pDel, accounting for ∼1% of total ASD cases, and nearly all carriers display some form of a NDD^37,39–41^. The 16pDel CNV mouse has been well-characterized with altered angiogenesis, corticogenesis, basal ganglia volume, ERK signaling, metabolism, and behavior^42–50^. Since individuals with this CNV have altered metabolic profiles, they may be particularly vulnerable to the biochemical changes following antibiotic-induced microbiome perturbations^51,52^.

Motivated by the link between early-life cephalosporin use and increased NDD risk, we investigated how a short-term therapeutic cefdinir course in early life affects gut microbiome composition, metabolite profiles, hippocampal development, and behavior in WT and 16pDel mice. Our findings reveal that this clinically relevant regimen perturbs the gut microbiome and induces NDD-related behavioral phenotypes. Notably, only cefdinir-exposed 16pDel mice showed altered hippocampal development, underscoring a gene-by-environment interaction in mediating vulnerability to early-life antibiotic exposure.

## Results

### A brief therapeutic exposure to cefdinir during early life perturbs the gut microbiome

Litters comprised of WT and 16pDel mice were randomly assigned to receive saline (control) or 10mg/kg cefdinir via gavage from postnatal (P) days 5-9 (**Fig1a**, with detailed sample sizes for all figures in **Supplementary file 1**). Microbial composition in the intestinal tract was profiled using 16S rRNA gene sequencing during the pre-weaning and weaning periods (P13 and P21, respectively). At P13, all cefdinir-exposed mice showed differences in microbial community structure (β-diversity) in the cecum compared to saline-exposed groups, indicated by both Jaccard and Bray-Curtis distances (**Fig1b, FigS1a**), with similar findings in the colon (**FigS1b-c**), and ileum (**FigS1d-e**). Although total bacterial abundances did not change (**Fig1c**), there also was reduced bacterial richness (α-diversity) in the cecum and colon (**Fig1d, FigS1f**), but not the ileum (**FigS1g**). Genus-level taxonomic shifts were assessed using the Microbiome Multivariate Association with Linear Models 2 (MaAsLin2) R package^53^, revealing reduced abundances of major taxa, including *Streptococcaceae*, *Lachnospiraceae* and *Ruminococcaceae*, alongside an increase in *Enterococcaceae* in the cecum of cefdinir- exposed mice (**Fig1e**). Similar results were observed in the colon and ileum (**FigS1h-i**). These microbial changes are unlikely to result from altered maternal care, as neither total body weight nor stomach weight differed between groups (**FigS1j-k**). By P21, α- diversity in cefdinir-exposed mice was no longer significantly different from saline controls (**Fig1f, FigS2a-b**), although β-diversity differences persisted within the cecum, colon, and ileum (**Fig1g, FigS2c-d**). Notably, several *Lachnospiraceae* and *Ruminococcaceae* genera continued to be reduced in cecal samples of P21 cefdinir- exposed mice (**Fig1h, FigS2e-f**), and these changes persisted in fecal specimens from P21 to P80 (**FigS3a-d**). Despite specific taxonomic changes, overall α- and β-diversity indices were comparable between saline- and cefdinir-exposed fecal specimens after P21 (**FigS4a-h**). Collectively, these results indicate that a brief early-life cefdinir exposure induces specific and persistent alterations in gut microbial composition, even as overall microbial diversity recovered during development.

**Figure 1:**
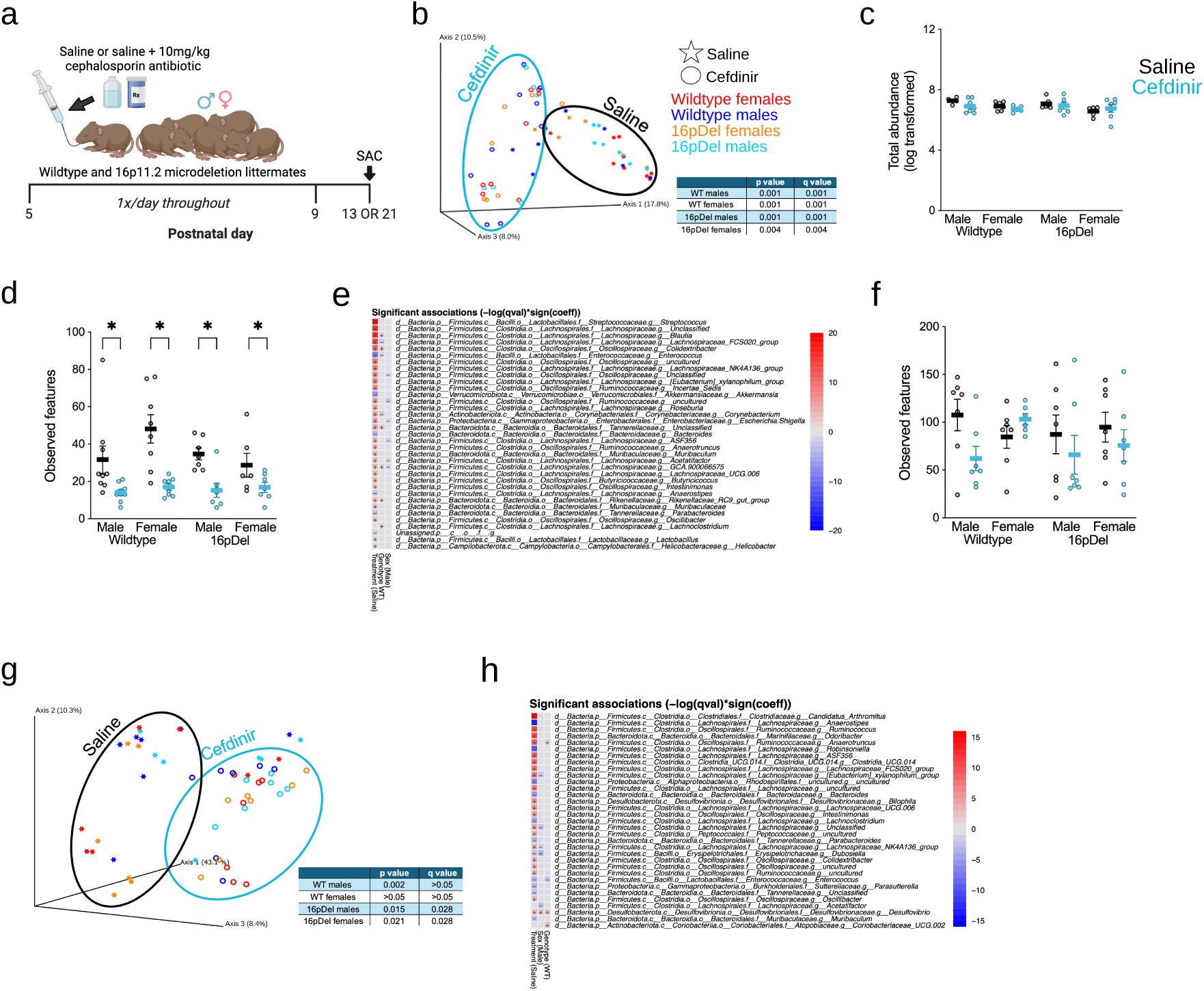
Early-life cefdinir exposure perturbed the gut microbiome. **a** Experimental design. Litters containing balanced ratios of WT and 16pDel pups were randomly assigned to receive saline or 10mg/kg cefdinir via oral gavage on P5-9. Offspring were sacrificed on P13 (panels b-e) or P21 (panels f-h) for experimentation and analysis of cecal samples. **b** Principal coordinate analysis (PCoA) plot showing microbial community composition based on Jaccard distances. **c** Log transformed total bacterial abundance. **d** Number of bacterial taxa at the genus level that were depleted following early-life cefdinir exposure. **e** Heatmap depicting abundances of the most variable taxa, classified by treatment, genotype, or sex. Depleted taxa are indicated in red, sustained taxa in blue, and the accompanying color bar reflects association strength. **f** Number of observed bacteria (genus level). **g** PCoA plot showing Jaccard distances. **h** Heatmap of the most variable taxa, analyzed by treatment, genotype, and sex. Group differences for PCoA plots (panels **b, g**) were assessed with permutational multivariate analysis of variance (PERMANOVA) pairwise tests using 999 permutations. Significance is indicated by *p* and *q* values <0.05. For taxa counts and abundances (panels **c-d, f**), comparisons were made using multiple Mann Whitney tests with false discovery rate (FDR) correction (two-stage step-up method of Benjamini, Krieger, and Yekutieli, *q* =1%). Data are presented as mean ± SEM, with \**q*<0.05 indicating significance. MaAsLin2 was used for differential abundance testing of log10-transformed taxon values (panels **e,h**). Supplementary file 1 for detailed sample sizes.

### Cefdinir-exposed mice display altered behavior throughout the lifespan

We next investigated whether mice with early-life microbiome perturbations exhibited behavioral phenotypes relevant to NDDs. In the absence of sex differences in the behavioral assessments, data from males and females were combined for analysis. Baseline genotype-specific behavioral traits in our 16pDel mice were confirmed by reduced exploratory behavior, decreased distance traveled, reduced time spent in the center of the arena, and increased time in the periphery of the open field arena at P60 (**FigS5a-d**)^46^. With this baseline established, we assessed the effects of cefdinir- induced microbiome perturbation on diverse behavioral outcomes (**Fig2a**). At P21, cefdinir-exposed mice showed reduced engagement with a novel sex- and age-matched mouse, compared to saline-exposed controls, an effect driven primarily by the 16pDel group (F (1, 107) = 4.31, *p*=0.040; Tukey’s adjusted *p*=0.049) (**Fig2b**). That all groups had similar levels of social odor detection and discrimination indicates that these social differences were not attributable to altered olfactory perception (**Fig2c**). At P80, cefdinir- exposed mice preferred interacting with a novel mouse over an object but did not show a preference for an unfamiliar mouse compared to the now familiar one from the prior trial, (F (1, 106) = 5.61, *p*=0.020) (**Fig2d, FigS5e**). Cefdinir-exposed mice also displayed reduced risk-aversion behavior, evidenced by increased time (F (1, 111) = 4.81, *p*=0.031) and entries (F (1, 111) = 6.36, *p*=0.013) into the open arms of the elevated plus maze at P63 (**Fig2e-f**). Lastly, cefdinir-exposed mice froze less in the 24hr contextual fear extinction trial, indicating compromised associative learning abilities, (F (1, 106) = 4.41, *p*=0.038) (**Fig2g**). No differences were observed in novel object recognition or the marble burying task across treatment or genotype (**FigS5f-g**). Collectively, these findings provide evidence that early-life cefdinir-induced microbiome perturbations can result in long-term, NDD-associated behavioral phenotypes persisting into adulthood.

**Figure 2:**
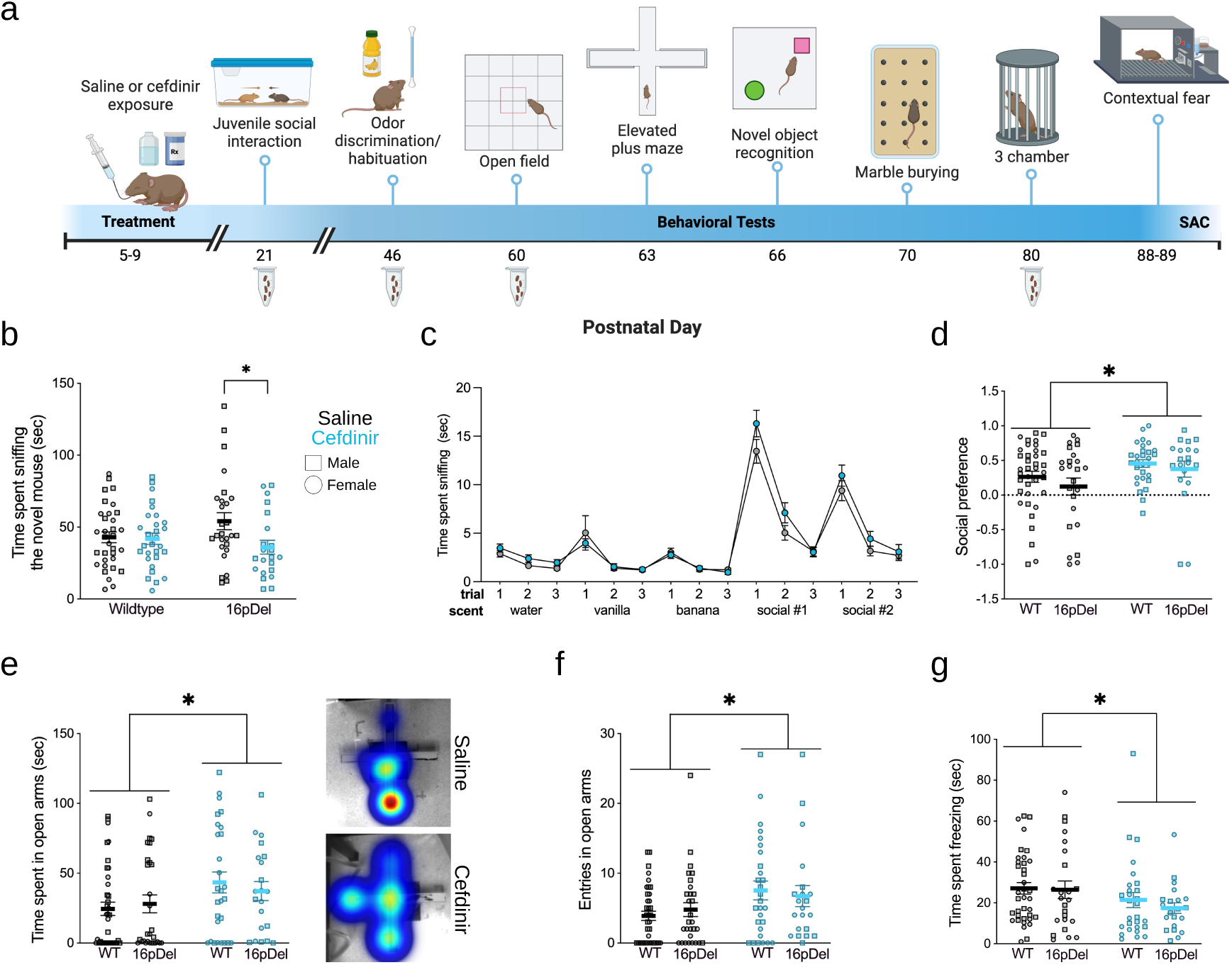
Cefdinir-exposed mice display altered behavioral outcomes throughout the lifespan. **a** Experimental paradigm. Litters were randomly assigned to receive saline or 10mg/kg cefdinir via oral gavage from P5-9. Offspring then underwent a longitudinal behavioral paradigm from P21-P89. Fecal pellets were collected at various timepoints for subsequent microbial analysis. **b** Social interaction assay comparing juvenile mice across treatment and genotype groups. **c** Social odor detection and discrimination assay assessing olfactory-based social recognition between saline- and cefdinir- exposed mice. **d** Social preference assay measuring time spent interacting with a novel mouse versus an inanimate object. The social preference index was calculated as the difference between time spent with a mouse and an inanimate object, divided by the total time spent with both; a positive score indicates a preference for the mouse, while a negative score indicates a preference for the object. **e-f** Elevated plus maze quantifying risk-aversion behaviors through time spent and entries into the open arms. **g** Contextual fear extinction assay measuring freezing behavior 24 hours after training. Comparisons in **b, d-g** were performed using two-way ANOVA; comparisons in **c** utilized two-way repeated measures ANOVA. Data are presented as mean ± SEM. See Supplementary file 1 for detailed sample sizes.

### Hippocampal alterations associated with an antibiotic-induced perturbed microbiome were exacerbated in 16pDel mice

To determine whether the cefdinir-induced behavioral changes were linked to altered neurodevelopment, we examined neurogenesis and astrogenesis in the hippocampus, a brain region critical for learning, memory, and behaviors associated with NDDs^33,54,55^.

Four days after antibiotic cessation (P13), cefdinir-exposed 16pDel males exhibited a robust decrease in the cell cycle regulator Cyclin E compared to saline-exposed controls while 16pDel females showed an increase (**Fig3a**). No antibiotic-induced differences were observed in WT male or female mice. Consistent with these results, cefdinir- exposed 16pDel males had parallel reductions in Ki67^+^ and EdU^+^ NSCs at P13, as well as a trend toward decreased Sox9^+^ astrocytes at P21 compared to saline-exposed controls (**Fig3b-d**). In contrast, compared with their controls, cefdinir-exposed 16pDel females showed no difference in Ki67^+^ or EdU^+^ NSCs at P13, but had a 53% increase in Prox1^+^ granule neurons at P21 (**Fig3b-c, e**). These alterations were not explained by blood brain barrier permeability, apoptosis, or direct antibiotic toxicity, as pilot studies revealed comparable findings between saline- and cefdinir-exposed mice (**FigS6a-d**).

**Figure 3:**
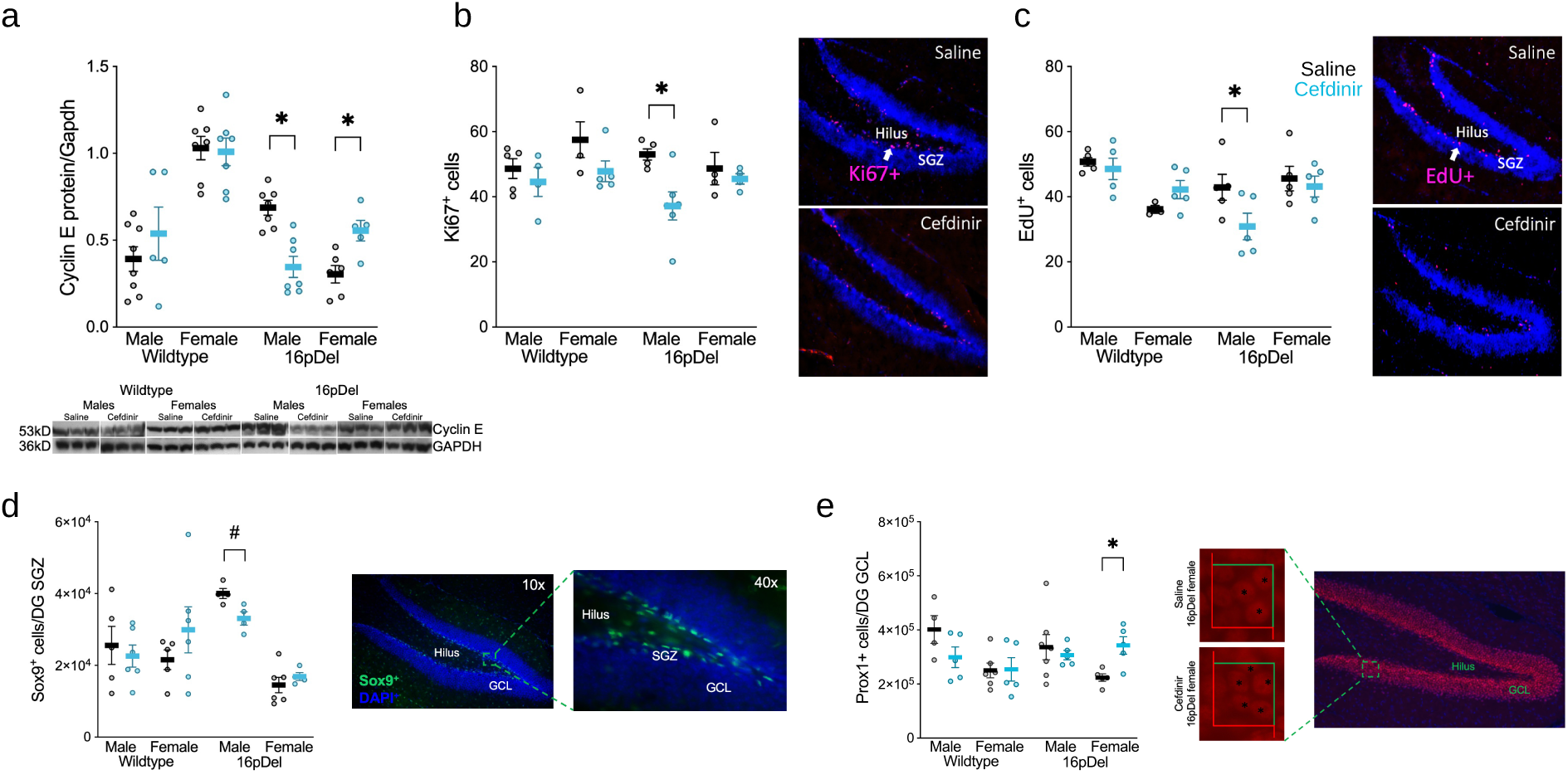
Cefdinir exposure alters genotype- and sex-dependent hippocampal development. **a** Western blot analysis of cyclin E protein normalized to GAPDH in individual 35ug hippocampal lysates from P13 mice. A total of 4 membranes were used, each containing 5-8 mice/treatment within the same genotype and sex. Representative bands from a subset of mice in each group are shown. **b-c** Quantification of proliferating cells in the P13 hippocampal dentate gyrus, measured by immunofluorescent labeling for **b** stem cell marker Ki67^+^ and **c** S-phase marker EdU^+^. Representative images from a saline- and cefdinir-exposed 16pDel male are shown. **d** Quantification of Sox9^+^ cells in the subgranular zone (SGZ) at P21. **e** Quantification of Prox1^+^ granule cell neurons in the granule cell layer (GCL) at P21. Representative images for **d** and **e** are provided. Comparisons for **a** were made using multiple Mann Whitney tests with the two-stage step-up method of Benjamini, Krieger, and Yekutieli, FDR *q*=1%, \**q*<0.05. Comparisons for **b-c** were made using a two-way ANOVA with Bonferroni’s multiple comparisons test, \**p*<0.05. Comparisons for stereology datasets **d-e** were made using multiple Mann Whitney tests with the two-stage step-up method of Benjamini, Krieger, and Yekutieli (FDR *q* =5%, \**q*<0.05, ^#^*q*<0.1). All data are presented as mean ± SEM. See Supplementary file 1 for detailed sample sizes.

Early-life cefdinir exposure also did not affect Iba1^+^ microglial cell numbers in the hilus at P21 (**FigS6e**). Overall, these findings demonstrate sexually dimorphic effects of early-life cefdinir-induced microbiome disruption on the hippocampus in 16pDel mice.

### Transcriptional changes in cefdinir-exposed 16pDel males overlap with autism- associated genes

To better understand the neurodevelopmental responses observed in the 16pDel mice, we examined hippocampal transcript levels to identify genes and pathways that varied in expression. As initial quality checks, we confirmed the appropriate expression of sex-determining genes in females and males, and the expected reduction of 16p11.2 interval gene expression in 16pDel mice (**FigS7a-d**). At P13, there were no significant differentially expressed genes (DEGs) between cefdinir- and saline-exposed mice when all genotypes and sexes were combined (**Fig4a**). Similarly, no DEGs were detected in the direct comparisons of saline- and cefdinir-exposed WT females or WT males (**Fig4b-c**). However, in 16pDel mice, compared to their saline-exposed controls, cefdinir exposure led to 5 DEGs in females and 118 DEGs in males (**Fig4d-e**). Notably, among the 118 DEGs in 16pDel males, 9 overlapped with curated lists of high-confidence ASD risk genes from the SFARI database^56^ and 4 from the ASC database^57^, with *Hdac4* and *Cadm1* present in both (**Fig4f**). These findings highlight transcriptional changes in 16pDel males induced by the early-life cefdinir exposure, including genes previously recognized as associated with high ASD risk in humans. At P21, 2 DEGs (*Lbhd2* and *Calcr*) between cefdinir- and saline-exposed mice were identified when all genotypes and sexes were combined (**Fig4g**). Subgroup analyses of direct saline- and cefdinir- exposed comparisons revealed substantial genotype- and sex-specific differences in gene expression: 130 DEGs in WT females, 190 in WT males, 22 in 16pDel females, and 261 in 16pDel males (**Fig4h-k**). Although DEGs were observed in each subgroup at P21, only 16pDel males (cefdinir- vs. saline-exposed) exhibited significant enrichment for gene ontology (GO) pathways (**Supplementary file 2**). Of the 108 upregulated DEGs in 16pDel males, terms related to myelination, glial cell development, and neurogenesis were overrepresented; of the 153 downregulated DEGs, terms related to axonogenesis, neurogenesis, cell-cell adhesion, and vesicle-mediated transport were enriched. The striking increase in both the number of DEGs and neurogenesis-related pathways from P13 to P21 indicates highly dynamic hippocampal transcriptional regulation across this developmental window. Notably, when comparing cefdinir- exposed 16pDel males to their saline-exposed counterparts at P21, >1,000 genes showed >three-fold expression differences. This increase highlights that cefdinir exposure in the 16pDel background leads to widespread dysregulation of hippocampal transcription during a critical developmental period. These differences were not confounded by litter effects, as each experimental sample represented a single mouse from a distinct litter. These DEGs also overlapped with ASD risk gene databases: 28 with SFARI, 18 with ASC, and 8 with both (**Fig4l**). Taken together, these expression data corroborate our protein and histological findings and demonstrate the early-life vulnerability of cefdinir-exposed 16pDel mice.

**Figure 4:**
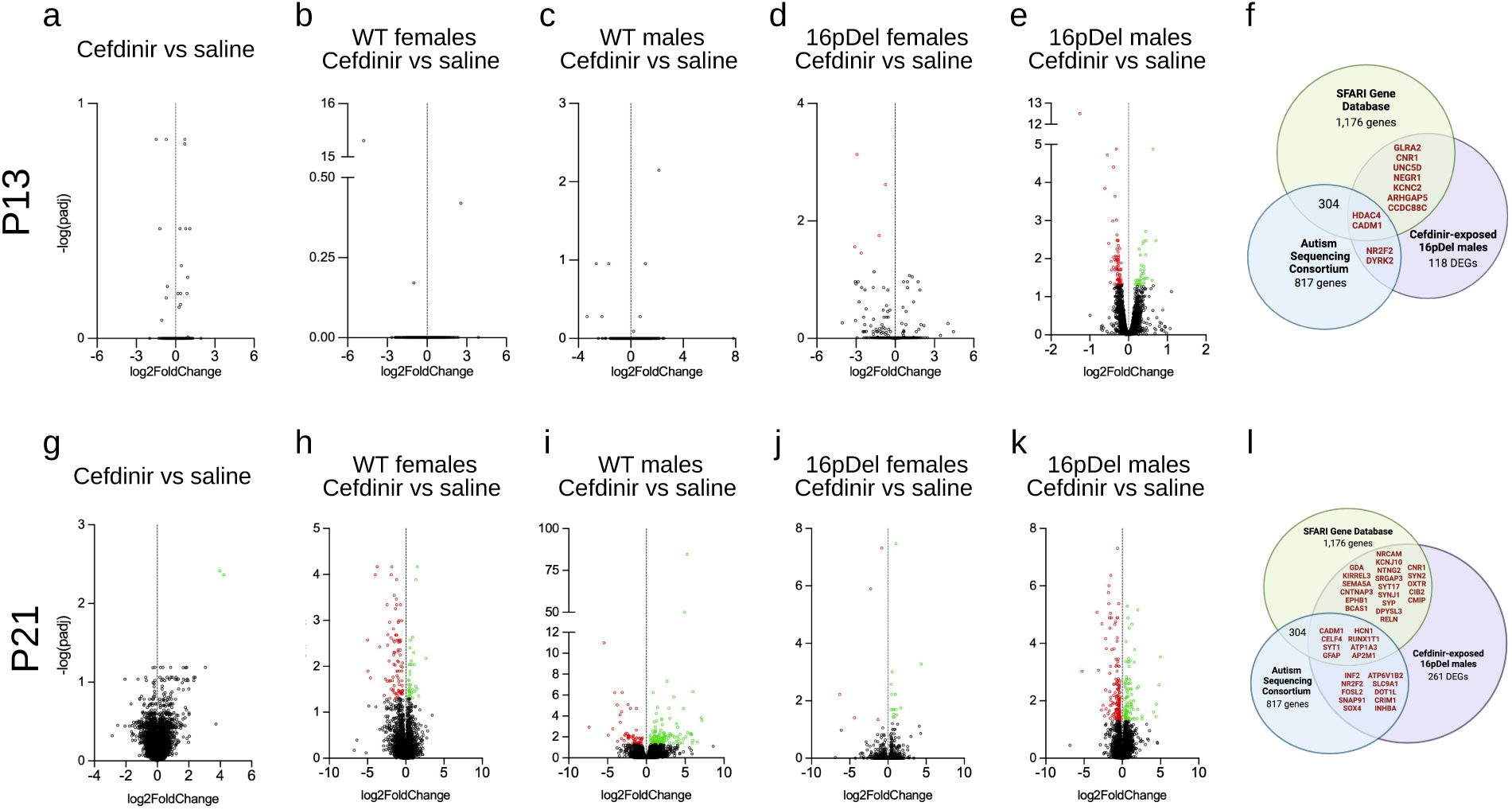
Differentially expressed genes in cefdinir-exposed 16pDel males overlap with human autism risk genes. Mice were sacrificed on P13 (panels a-f) or P21 (panels g-l), and one hippocampal hemisphere/mouse was randomly selected for bulk RNA sequencing. **a-e** Volcano plots depicting differentially expressed genes (DEGs) at P13 across **a** all cefdinir-exposed mice, or stratified by sex and genotype: **b** WT females **c** WT males **d** 16pDel females and **e** 16pDel males. **f** Venn diagram showing the overlap between 11 DEGs identified in cefdinir-exposed 16pDel males (purple) and ASD risk genes curated in the SFARI (green) and ASC (blue) databases. **g-k** Volcano plots showing DEGs at P21 for **g** all cefdinir-exposed mice or stratified by sex and genotype: **h** WT females **i** WT males **j** 16pDel females and **k** 16pDel males. **l** Venn diagram showing the overlap between 38 DEGs identified in cefdinir-exposed 16pDel males (purple) and ASD risk genes curated in the SFARI (green) and ASC (blue) databases. DEGs were detected using the DeSeq2 package in R. The Wald test was applied to generate p-values and log2fold changes. Given the exploratory design of this study and small sample size (n=3 mice/group, across 3 litters/treatment), significance was defined as an adjusted *p*<0.05. DEGs are depicted in green (upregulated), red (downregulated), and black [non-significant (padj>0.05)]. See Supplementary file 1 for detailed sample sizes.

### Cefdinir-exposed 16pDel mice display altered intestinal permeability, metabolites, and lipids

Given the neural and behavioral changes observed, we next explored whether cefdinir exposure altered gut barrier function and systemic metabolism, potentially linking microbiome perturbations to hippocampal alterations. We hypothesized that cefdinir exposure, in combination with the 16pDel risk variant, would impair intestinal barrier function (“leaky gut”), and subsequently affect lipid and metabolite profiles that could alter hippocampal development. We examined intestinal permeability and circulating metabolite and lipid profiles in both saline- and cefdinir-exposed mice at P11, midway between antibiotic cessation (P9) and the earliest neural changes observed (P13).

Compared to their saline-exposed counterparts, cefdinir-exposed 16pDel males showed a 3-fold increase in intestinal permeability, as indicated by elevated serum levels of orally gavaged FITC-dextran (F (1, 36) = 2.87, *p*=0.099; Tukey’s adjusted *p*=0.004); no changes were detected in other groups (**Fig5a-b**). Targeted serum metabolite analysis at P13 revealed that all cefdinir-exposed mice had reduced levels of pyruvate (F (1, 44) = 8.76, *p*=0.005), N,N-dimethylglycine (F (1, 44) = 8.93, *p*=0.005), cytosine (F (1, 44) = 4.22, *p*=0.046), 2-hydroxybutyrate (F (1, 44) = 7.01, *p*=0.011), and N-acetylglycine (F (1, 44) = 8.14, *p*=0.007) (**Fig5c-g**). Decreases in cholesterol ester (F (1, 43) = 6.60, *p*=0.014; Tukey’s adjusted *p*=0.046) and sterol (F (1, 43) = 8.44, *p*=0.006; Tukey’s adjusted *p*=0.027) content were observed exclusively in cefdinir-exposed 16pDel mice (**Fig5h-i**). To further explore the relevant metabolic pathways underlying these changes, we analyzed 29 metabolites correlated with cholesterol ester content in cefdinir-exposed 16pDel mice using the MetaboAnalyst KEGG database (**Fig5j**)^58,59^. This analysis identified alterations in arginine biosynthesis and glycerophospholipid metabolism, pathways with established roles in neural physiology and disease (**Fig5k**)^60–65^. By P21, most lipid differences had subsided; however, 6 new metabolites (5-hydroxylysine, nicotinamide adenine dinucleotide, glycine, sorbitol, creatinine, and proline) were reduced in all cefdinir-exposed mice, highlighting the dynamic nature of the metabolic effects of early-life cefdinir exposure (**Fig5l**).

**Figure 5:**
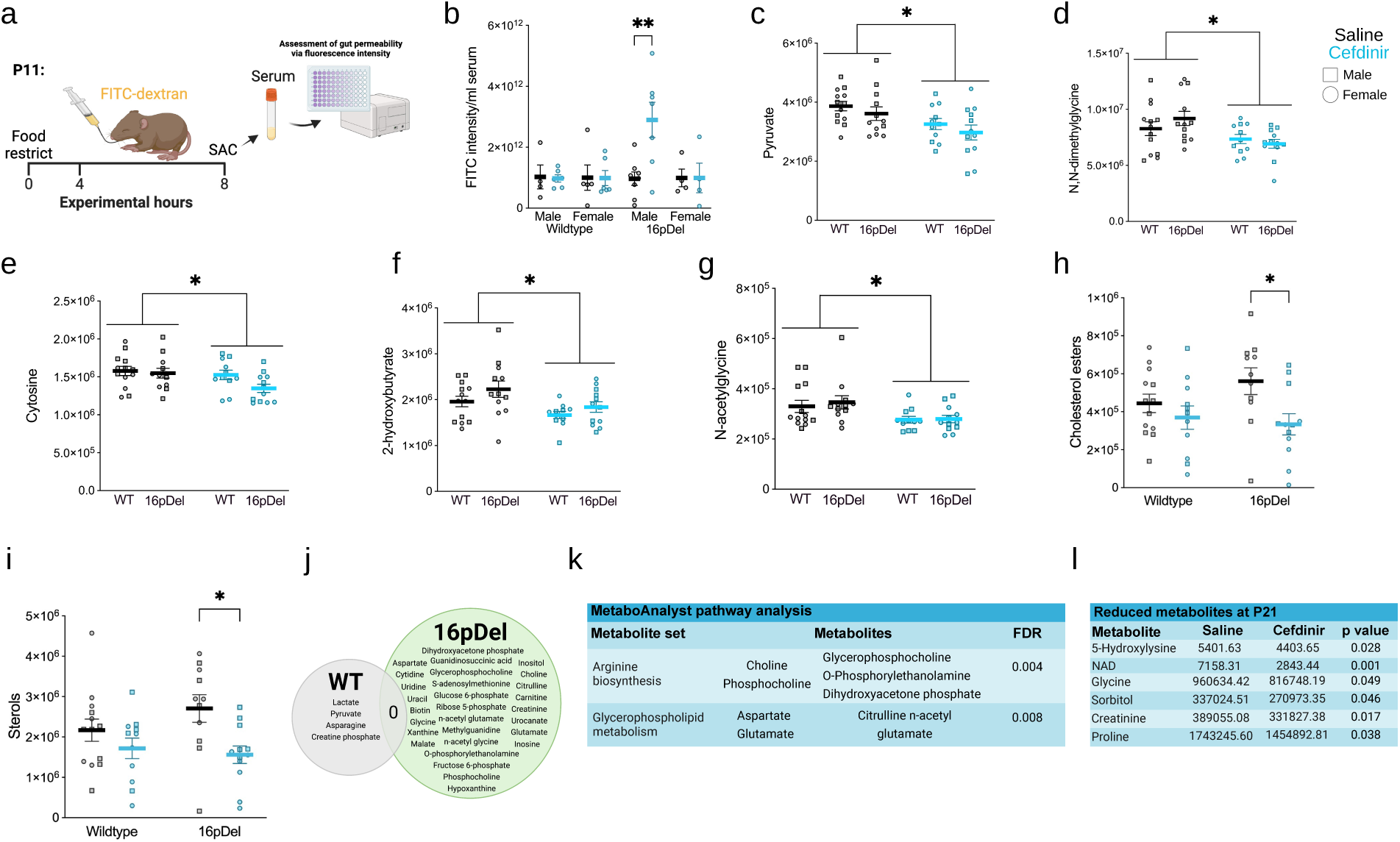
Distinct intestinal permeability and metabolic profiles in cefdinir- exposed 16pDel mice. **a** Experimental design for FITC-dextran assay. On P11, mice were fasted for 4 hours before receiving FITC-dextran by gavage. An additional 4 hours later, offspring were anesthetized and serum was collected for measurement of fluorescence intensity. **b** Measurement of serum FITC-dextran fluorescence intensity in P11 mice. **c-i** Peak intensity values of select metabolites and lipids at P13 assessed by targeted metabolomics. **j** Venn diagram displaying overlap of metabolites correlated with cholesterol ester content in cefdinir-exposed WT and 16pDel mice. **k** Table displaying enriched metabolite sets among 29 correlated metabolites in cefdinir- exposed 16pDel mice, based on the *Mus musculus* KEGG pathway library in MetaboAnalyst using the hypergeometric test for enrichment and relative-betweenness centrality as the topology measure. **l** Table summarizing metabolites reduced in cefdinir- exposed mice at P21. Comparisons for **b-i** were performed using a two-way ANOVA with Tukey’s multiple comparisons test, \*\**p*<0.01, \**p*<0.05. All data are shown as mean ± SEM. See Supplementary file 1 for detailed sample sizes.

Collectively, these findings reveal genotype-specific intestinal and metabolic responses that may interact with early-life cefdinir exposure to affect neurodevelopment.

## Discussion

Both clinical and epidemiological studies provide evidence associating antibiotic use in infancy with increased susceptibility to adverse health outcomes, including neuropsychiatric conditions^2–7,66–68^. We designed this preclinical study to model the therapeutic antibiotic courses commonly given to human infants; to define how gut microbiome alterations affect hippocampal development and behavior, and to determine whether host genotype modulates susceptibility. In our study, microbial alterations were most pronounced immediately after the antibiotic course (P13), with partial recovery by P21, as expected from a short antibiotic exposure. However, several taxa within *Lachnospiraceae* remained reduced throughout the lifespan of the cefdinir-exposed mice, consistent with reports in both antibiotic-exposed infants and adult rodents^67,69–71^. Depletion of *Lachnospiraceae* also has been linked to delayed development of the intestinal microbiome, disrupted metabolism, and altered neurodevelopment^67,69–71^.

Whether the observed behavioral phenotypes are driven by initial neurodevelopmental changes, the persistent *Lachnospiraceae* changes, or a combination of both remains to be determined.

Mice exposed to cefdinir during early life displayed NDD-related behavioral phenotypes across multiple domains that persisted for months after antibiotic cessation. Our findings are consistent with a prior study [Leclercq *et al*., (2017)], which examined the effects of a single antibiotic administered during early life^13^. In their model, low-dose penicillin V exposure from E12-P21 (4 weeks) resulted in decreased social preference and impaired risk-aversion behaviors at P42. Similarly, our 5-day postnatal therapeutic dose exposure also led to altered social behaviors, although the effects varied across measured modalities. Paralleling the findings of Leclercq et al. (2017), cefdinir-exposed mice spent more time in the open arms of the elevated plus maze at P60, reflecting reduced innate risk-avoidance behaviors^13^. By P89, nearly three months after initial antibiotic exposure, cefdinir-exposed mice demonstrated impaired extinction learning, a long-term deficit similar to findings observed in adult mice following prolonged treatment with a broad-spectrum antibiotic mixture^25^. Together, these consistent findings suggest that even brief antibiotic exposures can have lasting impacts on fear-aversion behaviors and learning, underscoring the importance of the gut microbiota in modulating neurobehavioral outcomes.

While cefdinir exposure induced changes in the gut microbiome and behavior across all groups, only 16pDel mice exhibited altered hippocampal stem cell and progenitor cell dynamics, supporting the hypothesis that NDD-related genetic variants exacerbate the neurodevelopmental impact of early-life antibiotic exposure. These effects were sexually dimorphic: males showed reduced NSC proliferation, while females had increased granule neuron numbers. Although microglial cell numbers did not differ between groups, further analyses of earlier ages, activation states and inflammatory markers are needed to fully understand immune responses. Cefdinir-related hippocampal transcriptional changes revealed a substantial gene-by-environment interaction, with the most extensive dysregulation and enrichment of ASD-linked pathways in 16pDel males. In these mice, DEGs overlapped with high-confidence ASD risk genes, demonstrating that host genotype influences the neurodevelopmental effects of early-life cefdinir exposure and its contribution to ASD risk. Kim *et al*., (2024) reported lower expression of ribosomal genes in the 16pDel males compared to wildtype controls, highlighting their inherent transcriptional differences^45^. We observed downregulation of ribosomal genes in 16pDel males after cefdinir exposure, indicating that environmental factors such as antibiotic treatment can accentuate baseline expression patterns. Notably, two of the DEGs we identified, *Hdac4* and *Cadm1*, both key regulators of cell cycle progression^63,64^ and present in major ASD gene databases^56,57^, were upregulated in this context. Elevated levels of *Hdac4* and *Cadm1*, each capable of inhibiting cell cycle progression, potentially contribute to the reduced NSC proliferation phenotype observed in cefdinir-exposed 16pDel males^72^.

To explore potential mechanisms underlying these neural changes, we assessed intestinal permeability and peripheral metabolic profiles. The increased intestinal permeability observed in cefdinir-exposed 16pDel males suggests another gene-by-environment effect, which may contribute to the neural alterations detected; future mechanistic studies are needed. The reductions in cholesterol esters and sterols in cefdinir-exposed 16pDel mice mirror clinical observations in some children with NDDs^73^, and may have neurodevelopmental significance, as cholesterol metabolites and sterols are critical for synapse formation, membrane integrity, and neurosteroid synthesis^74,75^.

In addition, metabolites involved in arginine biosynthesis and glycerophospholipid metabolism, uniquely altered in cefdinir-exposed 16pDel mice, may synergize with this CNV to affect hippocampal development^76,77^. The temporal specificity of these metabolic changes, which were present at P13 but largely resolved by P21, suggests that neurodevelopmental alterations may be established during a relatively brief but critical early window.

We note that recent comprehensive reviews have emphasized substantial conceptual and methodological limitations in studies linking gut microbiome perturbations to ASD^30^. Our findings are best interpreted as preliminary evidence for gene-by-environment interactions, rather than proof of causality. In summary, this exploratory study demonstrates that the 16pDel variant confers increased vulnerability to cefdinir-induced hippocampal dysregulation, with intermediate metabolic changes identified as possible mechanistic contributors. These data highlight the complex interplay between host genotype and environmental stressors in shaping neurodevelopmental outcome. This work also provides a foundation for future investigations of whether microbiome restoration following antibiotic exposures, through probiotics, prebiotics, or alternative modalities, can mitigate these neurodevelopmental abnormalities.

## Methods

### Ethical statement

All experimental protocols were conducted in compliance with NIH laboratory animal care guidelines and approved by the Rutgers University Institutional Animal Care and Use Committee (IACUC #PROTO999900063).

### Mouse husbandry

Heterozygous 16p11.2^+/-^ microdeletion (16pDel) male mice were purchased from The Jackson Laboratory (stock #013128; mixed B6/129 background). This line is maintained by breeding 16pDel males to B6129SF1/J “wildtype” (WT) females (stock #101043).

The resulting litters are comprised of ∼50:50 WT and 16pDel offspring. All mice were maintained on a 12hr light:12hr dark schedule where the lights were on from 6:00 a.m. to 6:00 p.m. Mice were housed in standard plastic cages in a colony room kept at ∼20°C, with ad libitum access to food and water.

### Genotyping

Mice were genotyped by PCR analysis of tail DNA, following standard protocols provided by Jackson Laboratories. Tail snips (0.5cm) were digested using a Extracta™ DNA Prep for PCR-Tissue (Quantabio). For 16pDel genotyping, the following primers were used: WT forward CCTGAGCCTCTTGAGTGTCC, WT reverse GTCGGTTCAGGTGGTAGACG, 16pDel forward ACCTGGTATTGAATGCTTGC, 16pDel reverse TGGTATCTCCATAAGACAGAATGC. These primers amplify a 194 bp band in both WT and 16pDel mice, and a 369-bp band in 16pDel mice only. Sex of the mice was confirmed using SRY primers: SRY forward TTGTCTAGAGAGCATGGAGGGCCATGTCAA and SRY reverse CCACTCCTCTGTGACACTTTAGCCCTCCGA, which yield a 273 bp band exclusively in male mice.

### Antibiotic Treatment

The cephalosporin antibiotic cefdinir (Alfa Aesar) was dissolved in 0.007% of 1N NaOH saline for administration. Control mice received the same 0.007% NaOH saline solution. Litters were randomly assigned to receive saline or 10mg/kg cefdinir via oral gavage once daily from P5-9. Prior to each gavage, mice were weighed to determine the appropriate dosing volume.

### Microbiome sample collection and processing

Offspring were sacrificed on P13 or P21, after which the cecum, colon, and ileum were collected, placed in cryopreservative tubes, snap frozen on dry ice, and stored at -80°C. Microbiota DNA was extracted using the PowerSoil-htp 96-Well DNA Isolation Kit (Qiagen). The V4 region of bacterial 16S rRNA genes was amplified in triplicate reactions using barcoded fusion primers 515F/806R, which amplifies bacterial and archaeal 16S genes. Amplicon concentrations were quantified for each sample using the Quant-iT PicoGreen dsDNA assay kit (Life Technologies), and samples were pooled in equal amounts. These pooled amplicons were purified with the Qiaquick PCR purification kit (Qiagen) to remove primers, quantified using the high-sensitivity dsDNA assay kit and Qubit 2.0 Fluorometer (Life Technologies), then combined at equal concentrations to create the sequencing library. The ∼254 bp V4 amplicons were sequenced with the Ilumina MiSeq 2x150 bp platform at Azenta Life Sciences (South Plainfield, NJ). Fecal pellet samples from mice used in behavioral studies were processed similarly at P21, P45, P60, and P80.

### 16S rRNA sequencing analysis

Quality filtering and downstream analysis were performed using quantitative insights for microbial ecology 2 (QIIME)^78^. Sequence reads were trimmed, denoised, merged, and chimeric sequences were removed using DADA2^79^ to generate a high-resolution feature table. α-diversity was quantified using the number of observed features, while β- diversity was assessed using both Jaccard and Bray Curtis matrices^80,81^; differences in β-diversity were visualized with principal coordinate analysis plots. Taxonomic classification was performed using the SILVA reference database^82^. Differentially abundant bacterial taxa were identified with the Microbiome multivariate association with linear models (MaAsLin2) algorithm^53^. These analyses included measures of α- diversity, β-diversity, and community composition.

### Immunoblotting

Equal amounts of protein (35μg/lane) from either P13 or P21 hippocampal lysates, or P11 whole brain lysates, were loaded on 12% acrylamide gels. Separated protein in gels were transferred to membranes, which were then blocked in 5% milk prepared with 0.05% Triton X-100 in 1X TBS. Membranes were incubated with primary antibody overnight at 4°C. After washing, HRP-conjugated secondary antibody (1:1,000; NA931V or NA934V, GE Healthcare) were applied for 1hr at room temperature (RT). Protein bands were visualized using chemiluminescence (ECL; GE Healthcare). Primary antibody dilutions were: anti-Cyclin E (1:300; sc-377100, Santa Cruz), anti-Cleaved Caspase-3 (1:1,000; 9664S, Cell Signaling) anti-Occludin (1:1,000; 40-4700, Abcam) and anti-GAPDH (1:10,000; 60004-1-Ig, Proteintech). Band intensities were normalized to GAPDH and quantified by densitometry.

### EdU administration and cryopreservation

In separate studies, P6 and P13 offspring received a subcutaneous injection of 100mg/kg 5-ethynyl-2’-deoxyuridine (EdU; E10187, Invitrogen) to assess precursor S-phase entry. EdU solutions were freshly prepared on the day of injection by solubilization in 0.007% 1N NaOH. Two hours after injection, mice were anesthetized and transcardially perfused with 1X PBS, followed by ice cold, freshly prepared 4% PFA (pH 7.40). Brains were post-fixed in 4% PFA for 24hrs at 4°C, cryoprotected in 30% sucrose, embedded in Tissue-Tek (Sakura), and stored at -80°C until cryosectioning.

### Brain tissue processing

The left hemispheres of P6 and P13 mice were sectioned at 14μm thickness in the sagittal plane using a cryostat (Leica, Heidelberg, Germany). Every other section was collected and mounted onto charged, uncoated Superfrost slides (VWR). For P21 stereological analyses, brains were cut into 30μm thick sagittal sections on a cryostat in a 1:10 series fashion.

### Immunohistochemistry

Slides were first rinsed with 1X PBS to remove residual OCT. Antigen retrieval was then performed by incubating sections in boiling 1X citrate buffer for 10mins at 100°C, followed by gradual cooling to 82-85°C for an additional 10mins. Sections were subsequently blocked in 33% normal goat serum (NGS) with 0.5% Triton X-100 in 1X PBS for 1hr at RT and incubated overnight at 4°C with primary antibody prepared in 2% NGS with 0.3% Triton X-100 in 1X PBS. The following day, sections were washed with 1X PBS and incubated with secondary antibody for 1hr at RT. Sections were then counterstained with DAPI (1:1,000; D21490, Invitrogen) in 1X PBS at RT for 10mins, washed in 1X PBS, and coverslipped. Primary antibodies include anti-Ki67 (1:200; 5556003, BD Pharmigen), anti-Prox1 (1:1,000; AB5475, Millipore Sigma), anti-Sox9 (1:1,000; ab185966, Abcam), and anti-Iba1 (1:1,000; ab5076, Abcam), and secondary antibodies include goat anti-mouse 594 or goat anti-rabbit 488 (1:500; A11005 or A11008, Invitrogen).

### Click-iT EdU

Slides were first rinsed with 1X PBS to remove residual OCT. Sections were permeabilized with 0.5% Triton X-100 in 1X PBS for 10mins and then washed twice with 3% BSA in 1X PBS for 5 mins/wash. The EdU staining solution was prepared (for 1mL solution): 860µl 1x Tris-HCl (pH 8.6), 40µl 100mM CuSO4 (stock 16mg/ml in H2O; C495- 500, Fisher Scientific), 2.5µl Alexa fluor azide (A10270, Thermo Fisher Scientific) and 100µl reaction buffer additive (Ascorbic acid, 200mg/ml H2O; A92902, Millipore Sigma). Sections were incubated in the EdU staining solution for 30mins at RT, protected from light. After incubation, sections were washed once with 3% BSA in 1X PBS, counterstained with DAPI (1:1,000) for 10mins, rinsed with 1X PBS, and then coverslipped.

### Cell quantification

Sections were imaged using an Axiovert 200M microscope (Zeiss, USA). Total numbers of EdU^+^ and Ki67^+^ cells in P6 and P13 offspring were quantified in the hilus and subgranular zone of the hippocampal dentate gyrus on 40X and 20X, respectively. For P13 analyses, cell counts were obtained from an average of 8 sections/mouse. For P6 16pDel males, quantification was performed from ∼6 sections/mouse. Iba1^+^ cells in P21 offspring were counted within the hilus at 40x magnification, using ∼8 sections/mouse.

All analyses were performed by an experimenter blinded to treatment, sex, and genotype.

### Stereological analysis

Sections were imaged on a ZEISS Axio Imager M2 microscope equipped with Stereo Investigator (MBF Bioscience). Quantification was performed using the optical fractionator workflow, with sampling parameters set to a counting frame of x:15 y:15, a grid size of x:47 y:47, and 10% sampling fraction for each outlined region. Brains with fewer than five viable sections were excluded from analyses. Sections were not used if excessively lateral or medial, damaged, folded, or lacked the dentate gyrus. The z-axis thickness of each section was measured and recorded for population estimates. Only datasets with a Gunderson coefficient of error (m=1) <0.1 were included. Quantifications were performed blind to genotype, sex, and treatment in every 10th section through the entire hippocampus. Total Prox1^+^ cells were quantified within the granule cell layer, with regions traced at 10x magnification and nuclei counted at 63x oil immersion. Sox9^+^ cells were quantified in the subgranular zone, and Iba1^+^ cells were quantified in the hilus.

Analyses were conducted on ∼5-9 sections/mouse.

### Intestinal permeability

The intestinal permeability assay was performed as described in Hsiao et al., (2013)^10^. On P11, offspring were fasted for 4hrs prior to receiving an oral gavage of 0.6g/kg 4- kDa FITC-dextran (46944, Millipore Sigma). Four hours after gavage, offspring were anesthetized, and serum was collected and assessed for fluorescence intensity (485 excitation, 528 emission) on the SpectraMax iD3 (Molecular Devices). Standard curves were generated using 1:2 serial dilutions from 0-12.8mg/mL in saline. Serum samples from a non-FITC gavaged mouse (control) and all experimental mice were diluted 1:20 in saline, and all standards and samples were run in triplicate on 96-well plates.

### Serum lipidomics and metabolomics

Blood was collected from mice in the P13 and P21 microbiome studies and centrifuged at 5,000 RPM for 10mins at 4°C to isolate serum, which was stored at -80°C until analysis. Serum samples were extracted for both lipids and polar metabolites and analyzed using positive and negative ionization modes at the Metabolomics Shared Resource (MSR) of the Rutgers Cancer Institute of NJ^83–87^. Lipid identification was performed with MS-DIAL, using in silico database annotations. Metabolic profiling was conducted in an untargeted fashion on a Q Exactive Quadrupole-Orbitrap mass spectrometer coupled to Vanquish Horizon UHPLC system. The MSR in-house metabolite database includes over 400 polar metabolites, each with established retention times and accurate mass measurements.

### Bulk RNA sequencing

The hippocampus was dissected from a subset of mice used in the P13 and P21 microbiome and metabolome studies. For each sample, one hemisphere was randomly assigned for protein analysis, while the other was immediately stored in RNAlater (Thermofisher) at 4°C for 24hrs, then the solution was removed and the tissue stored at -80°C. Dissections were performed using micro forceps and a dissecting microscope, referencing anatomical landmarks from the Allen Institute Developing Mouse Brain Atlas^88^. For gene expression studies, RNA isolation, quality control, and mRNA sequencing was performed by Azenta Life Sciences. RNA integrity number values ranged from 6.1-9.6. Libraries were sequenced on the IluminaHiSeq 2x150 bp platform. Sequence reads were trimmed using Trimmomatic v.0.36^89^, followed by alignment to the *Mus musculus* GRCm38 reference genome (ENSEMBL) using STAR v.2.5.2b^90^.

Unique gene hit counts within exon regions were quantified with featureCounts (Subread package v.1.5.2)^91^.

### Juvenile social interaction

Juvenile reciprocal social interactions were assessed on P21, immediately prior to weaning. Testing was conducted in a sterile home cage containing a 0.5cm layer of clean bedding. Each subject mouse was singly housed in a clean cage for 1hr before the test. Following this brief isolation, the subject mouse and an age- and sex-matched C57BL/6J partner were simultaneously placed in the arena and their interactions were videorecorded for 10mins, as described in Portmann et al., (2014)^43^. Videos were analyzed by observers blinded to treatment, genotype, and sex. Parameters of social behaviors included nose-to-nose sniff, front approach, and nose-to-anogenital sniff. All behaviors were analyzed for frequency of occurrence.

### Odor habituation/discrimination task

On P45, experimental mice were individually placed into clean home cages and isolated for 1hr. Following isolation, each mouse was exposed to a series of odors presented on saturated cotton swabs, following the protocol adapted from Arbuckle et al., (2015)^92^.

The task consisted of 15 consecutive 1min trials: 3 trials each with tap water, 1:100 diluted vanilla extract (McCormick®), 1:100 diluted banana extract (McCormick®), social odor from 129/SvlmJ mice, and social odor from C57BL/6J mice. Social odor stimuli were prepared by wiping a swab in a zigzag motion across the bedding of a soiled cage containing unfamiliar mice of the same sex. Sniffing behavior was recorded live by an observer positioned ∼2m from the testing cage, blinded to treatment and genotype.

Sniffing was scored when the mouse’s nose was within 1cm of the odor swab, and the total time spent sniffing was measured for each trial.

### Open field exploration

On P60, mice were placed in a clean, novel arena for 30mins under dim lighting conditions (100 lux). Locomotor activity and spatial location were recorded using infrared beam breaks and analyzed with Kinder Scientific software. Behavioral parameters, including rearing events, total distance traveled, number of entries, and time in each compartment (center, periphery, and total) were quantified.

### Elevated plus maze

On P63, mice were tested in the elevated plus maze task to assess risk-avoidance behavior. Each mouse was placed in the center of the maze and allowed to freely explore the apparatus for 10mins. All sessions were videorecorded and analyzed using Ethovision XT software (Noldus). Behavioral measures included distance traveled, time spent, and number of entries into both open and closed arms of the maze.

### Novel object recognition

On P66, mice underwent three 10min trials following a protocol adapted from Leger et al., (2013)^93^. The first trial (habituation) involved free exploration of an empty arena. In the second trial, mice were exposed to two identical inanimate objects. Ninety minutes later, in the third trial, one of the previously presented objects (familiar) was replaced with a novel object to assess recognition memory. All sessions were videorecorded and tracked using EthoVision. Object exploration was defined as time spent sniffing the object with the nose oriented toward it at a distance of less than 2cm. Recognition memory was defined by significantly greater time spent sniffing the novel object relative to the familiar one.

### Marble burying

On P70, each mouse was placed individually into a clean home cage containing ∼4cm of corncob bedding and ten evenly spaced blue glass marbles arranged in five rows. Mice were allowed to explore the cage freely for 30mins. At the end of the session, photographs of each cage were taken to document marble displacement and burial. To reduce subjectivity in quantification, marble burying was analyzed using a modified ImageJ script based on Wahl et al., (2022)^94^.

### Three-chamber sociability task

Social preference and social novelty were assessed using a three-chamber task protocol adapted from Rein et al. (2020)^95^. On P80, each mouse completed three consecutive 10min trials: (1) a pre-test habituation to the empty chamber and wire cups; (2) a social preference trial, in which one cup contained an inanimate object and the other an unfamiliar, sex- and aged- matched mouse; and (3) a social novelty trial, in which the mouse from trial 2 (now familiar) remained, while a new mouse replaced the inanimate object. All sessions were videorecorded and tracked using EthoVision.

### Contextual fear conditioning

Contextual fear conditioning was performed in a Freezeframe Chamber (Coulbourn Instrument) following the protocol described by Hagewoud et al., (2010)^96^. The context included a light, fan, and peppermint odor as cues. On P88, mice were fear conditioned, and freezing behavior was assessed 24hrs later on P89. During conditioning, mice were allowed to explore the chamber for 3mins before receiving a 2sec, 0.70mA foot shock, after which they remained in the context for an additional 15sec before being returned to their home cages. To test associative learning, mice were re-exposed to the same chamber 24hrs later, and the percentage of time spent freezing during a 3min session was measured. All behavioral scoring was performed blind to treatment, genotype, and sex.

### Statistical analysis

#### Microbiome composition

For microbiota studies, significant differences in α-diversity between experimental groups were determined using multiple Mann-Whitney tests, with a false discovery rate (FDR) correction (*q* =1%) performed by the two-stage step-up method of Benjamini, Krieger, and Yekutieli^97,98^. β-diversity differences were assessed using permutational multivariate analysis of variance (PERMANOVA) pairwise tests with 999 permutations^99^. Statistical significance was defined as *p* and *q*<0.05. Differences in the relative abundance of taxa were evaluated using the MaAsLin2 R package^53^, with significance thresholds set at the default parameters.

#### Behavior

Datasets were analyzed using two-way ANOVA followed by Tukey’s multiple comparisons test, except for the odor discrimination/habituation task, which was analyzed using a two-way repeated measures ANOVA^100,101^. Significance was set at *p*<0.05.

#### Immunoblotting

Each gel was restricted to 14 lanes after accounting for the protein ladder. The primary objective was to evaluate treatment-dependent effects. Therefore, significant differences between treatment groups (saline vs cefdinir) within sex and genotype were assessed using multiple Mann Whitney tests, with FDR correction *q* =1% applied via the two-stage step-up method of Benjamini, Krieger, and Yekutieli^97,98^.

#### Immunohistochemical quantification

EdU and Ki67 quantification data were analyzed using two-way ANOVA followed by Bonferroni’s multiple comparisons test, with significance set at *p*<0.05^100,102^. Stereological data were analyzed using multiple Mann-Whitney tests with FDR correction *q* =5% applied via the two-stage step-up method of Benjamini, Krieger, and Yekutieli^97,98^.

#### Bulk RNAseq

Gene expression comparisons were analyzed using the DESeq2 R package^103^. The Wald test was applied to estimate *p* values and log2fold changes for each gene^104^. Genes with an adjusted *p*<0.05 were considered differentially expressed genes in each comparison.

#### Intestinal permeability

All standards and samples were assayed in triplicate on a 96-well plate. Data were analyzed using a two-way ANOVA followed by Tukey’s multiple comparisons test, with statistical significance set at *p*<0.05^100,101^.

#### Sera lipidome and metabolome

Datasets were analyzed using ordinary two-way ANOVA with Tukey’s multiple comparisons test, with significance set at *p*<0.05^100,101^.

#### Correlations and pathway analysis

Pearson correlations and linear regressions were performed between the metabolic and lipid datasets^105,106^. Correlations with *p*<0.05 were uploaded to MetaboAnalyst for pathway analysis using the *Mus musculus* KEGG library^58,59^. Enrichment analysis was conducted with the hypergeometric test, and pathway topology was assessed using relative-betweenness centrality as the topological measure^107^. Metabolite sets were considered significantly enriched at FDR<0.05.

### Respective contributions

CRM, JHM, BAS, MJB, and EDB contributed to conceptualization of the study. CRM performed oral gavage on all mice used for experimentation. CRM, ZG, XZhou, and XZhang dissected brain and gastrointestinal tissues for processing. CRM collected serum for metabolomic and lipid studies. CRM and AH performed the FITC-dextran assay. CRM performed immunoblotting studies. CRM, AM, CN, KK, DT, DP, ZM, and AH performed immunohistochemistry and histology quantifications. CRM, AM, KK, DT, CN, DP, ZM performed and analyzed behavioral studies. CRM, ZG and JX contributed to data analysis. CRM wrote the manuscript, EDB revised initial drafts, EDB, MJB, JHM, BAS, JX, JA, and STV revised the final version.

## Notes

### Competing Interest Statement

The authors have declared no competing interest.

### Summary of Updates

The revised title more concisely emphasizes the gene-by-environment effects observed in this study. The experimental design, datasets, and statistical outcomes are unchanged, but the results text was edited to clarify effect sizes, sex/genotype specificity, and the dynamic nature of these gene-by-environment-dependent responses. The discussion was revised to acknowledge limitations and the absence of a causal mechanism, place the work within critiques of microbiome ASD studies, deepen mechanistic interpretation (leaky gut, cholesterol/sterol changes, arginine and glycerophospholipid pathways), and propose specific future experiments (cell cycle kinetics, microglial activation and cytokines, gut barrier assays, and microbiome restoration approaches).

