## Supplemental Figures for "The 16p11.2 microdeletion exacerbates neurodevelopmental alterations induced by early-life microbiome perturbation"

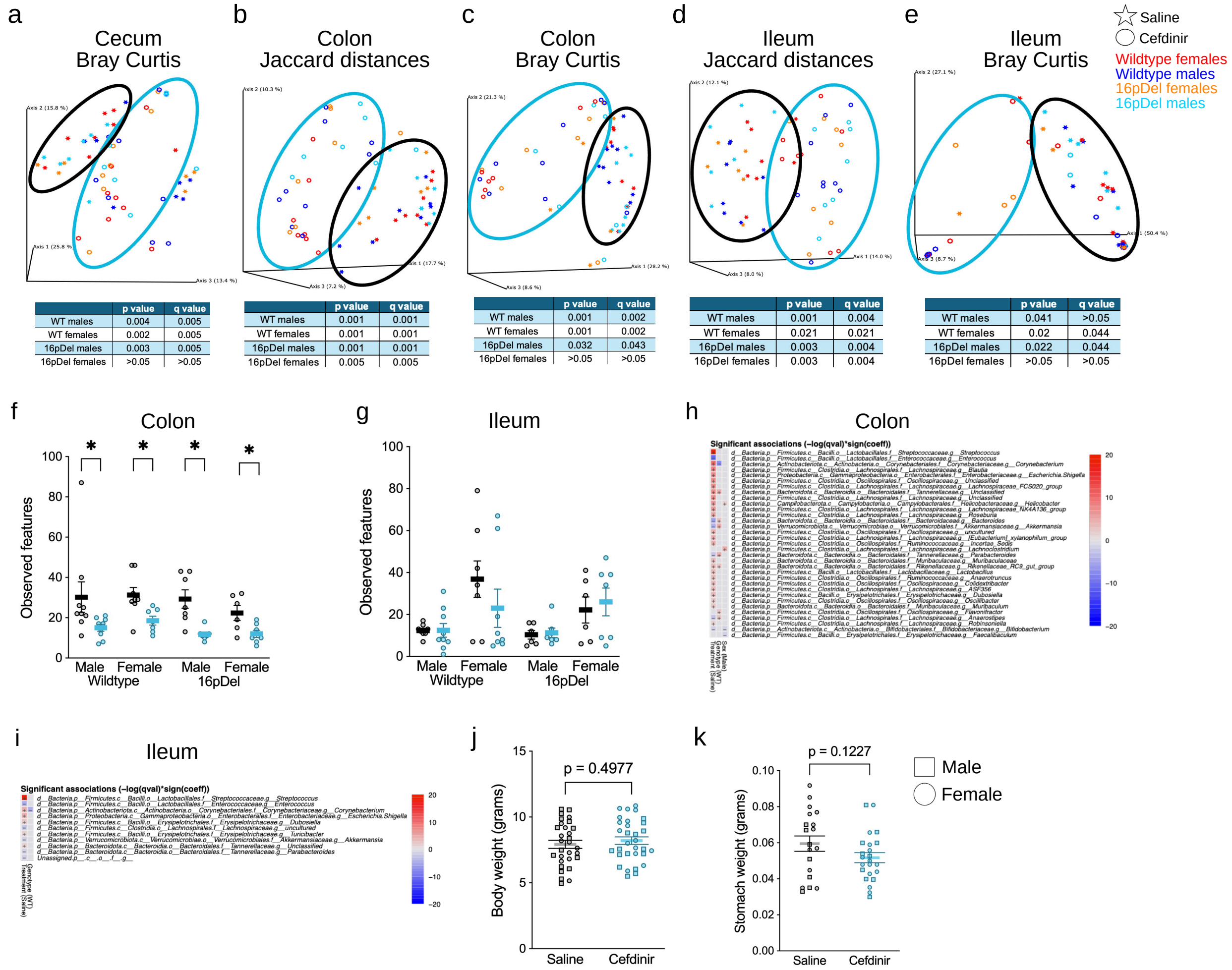


a (Cecum, Bray Curtis)

| | p value | q value |
|---|---|---|
| WT males | 0.004 | 0.005 |
| WT females | 0.002 | 0.005 |
| 16pDel males | 0.003 | 0.005 |
| 16pDel females | >0.05 | >0.05 |

b (Colon, Jaccard distances)

| | p value | q value |
|---|---|---|
| WT males | 0.001 | 0.001 |
| WT females | 0.001 | 0.001 |
| 16pDel males | 0.001 | 0.001 |
| 16pDel females | 0.005 | 0.005 |

c (Colon, Bray Curtis)

| | p value | q value |
|---|---|---|
| WT males | 0.001 | 0.002 |
| WT females | 0.001 | 0.002 |
| 16pDel males | 0.032 | 0.043 |
| 16pDel females | >0.05 | >0.05 |

d (Ileum, Jaccard distances)

| | p value | q value |
|---|---|---|
| WT males | 0.001 | 0.004 |
| WT females | 0.021 | 0.021 |
| 16pDel males | 0.003 | 0.004 |
| 16pDel females | 0.003 | 0.004 |

e (Ileum, Bray Curtis)

| | p value | q value |
|---|---|---|
| WT males | 0.041 | >0.05 |
| WT females | 0.02 | 0.044 |
| 16pDel males | 0.022 | 0.044 |
| 16pDel females | >0.05 | >0.05 |

**Supplemental figure 1 (related to figure 1). Microbial changes in the colon and ileum of P13 mice following early-life cefdinir exposure.**

**a-e** Principal coordinate analysis (PCoA) plots of Bray Curtis or Jaccard distances, displaying the composition of cecal, colon, and ileum microbiota samples. **f-g** Genus-level bacterial diversity in **f** colon and **g** ileum samples. **h-i** Heatmaps illustrating the most variable taxa, classified by treatment, genotype, and sex in the **h** colon and **i** ileum, with red indicating depletion and blue indicating sustained bacteria; color intensity reflects association strength. **j** Total body weight measurements at P13. **k** Milk-containing stomach tissue weights at P11. Comparisons for **a-e** were performed using PERMANOVA pairwise tests with 999 permutations. Significant differences were inferred with *p* and *q* values <0.05, provided in the corresponding tables. Comparisons for **f-g** were made using multiple Mann Whitney tests with the two-stage step-up method of Benjamini, Krieger, and Yekutieli false discovery rate (FDR) *q* =1%. Data are presented as mean ± SEM, *q <0.05. Significance for **h-i** was inferred by MaAsLin2 analysis using Log10 taxon. Comparisons for **j-k** were made using the Student’s t-test. Detailed sample sizes are reported in Supplementary file 1.

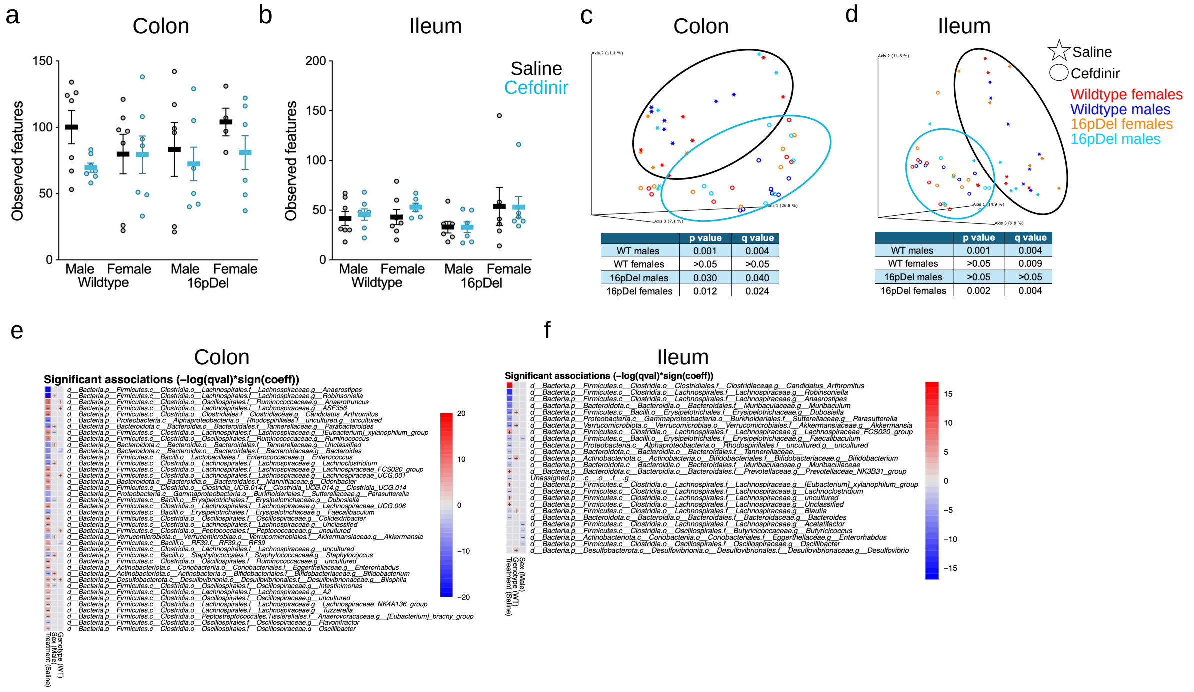


**Supplemental figure 2 (related to figure 1). Microbial changes in the colon and ileum of P21 mice following early-life cefdinir exposure.**

**a-b** Analysis of $\alpha$-diversity in **a** colon and **b** ileum samples. **c-d** PCoA plots depicting Jaccard distances for individual **c** colon and **d** ileum microbiota samples. **e-f** Heatmaps displaying the top taxa that differed in relative abundances according to treatment, genotype, and sex in the **e** colon and **f** ileum. Red indicates depleted bacteria and blue indicates sustained bacteria, with color intensity reflecting the strength of association. Comparisons for **a-b** were made using multiple Mann Whitney tests with the two-stage step-up method of Benjamini, Krieger, and Yekutieli (FDR $q$ =1%), with data presented as mean ± SEM. Comparisons for **c-d** were assessed using PERMANOVA pairwise tests with 999 permutations; significant $p$ and $q$ values <0.05 are provided in the tables. Differential taxa for **e-f** were identified using MaAsLin2 analysis on Log10 taxon. Detailed sample sizes are reported in Supplementary file 1.

a

P21

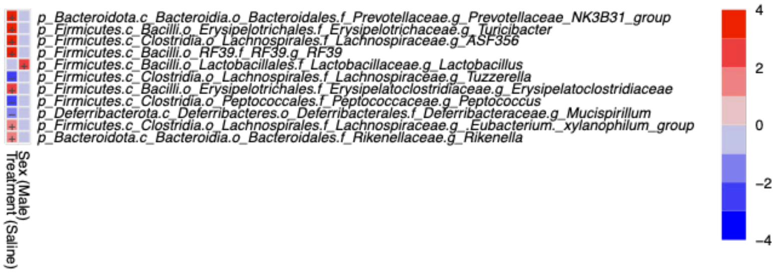


b

P45

| Taxon | Group | Coefficient | pval | qval |
|---|---|---|---|---|
| p_Firmicutes.c_Clostridia.o_Clostridia_UCG.014.f_Clostridia_UCG.014.g_Clostridia_UCG.014 | Saline | 0.371 | 0.0004 | 0.101 |
| p_Firmicutes.c_Clostridia.o_Lachnospirales.f_Lachnospiraceae.g_A2 | Saline | 0.554 | 0.0022 | 0.132 |
| p_Bacteroidota.c_Bacteroidia.o_Bacteroidales.f_Rikenellaceae.g_Rikenella | Saline | 0.434 | 0.0021 | 0.132 |
| p_Firmicutes.c_Bacilli.o_RF39.f_RF39.g_RF39 | Saline | 0.272 | 0.0011 | 0.132 |
| p_Firmicutes.c_Clostridia.o_Lachnospirales.f_Lachnospiraceae.g_ASF356 | Saline | 0.546 | 0.0032 | 0.157 |

c

P60

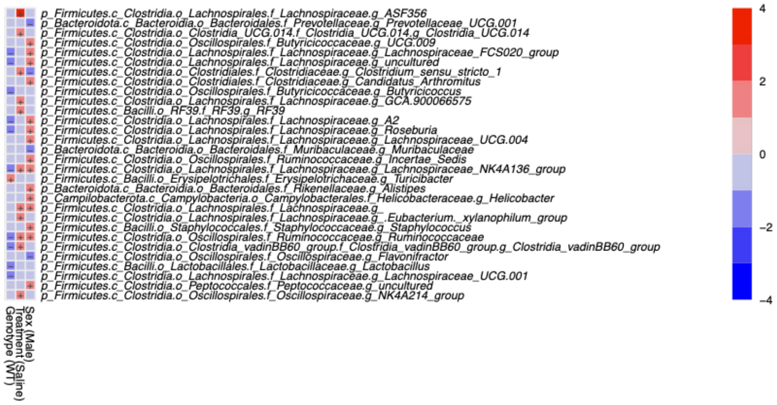


d

P80

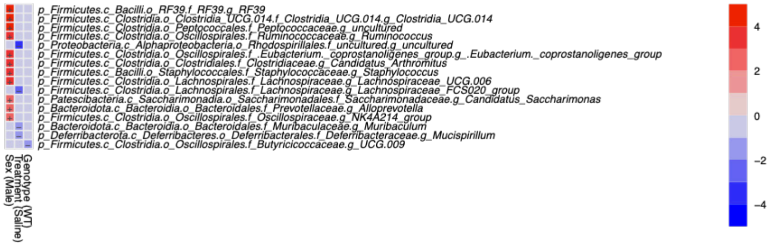


**Supplemental figure 3 (related to figure 1). *Lachnospiraceae* and *Ruminococcaceae* abundances in fecal samples are persistently shifted throughout the lifespan**.

**a-d** Heatmaps displaying the most variable taxa in fecal samples, classified by treatment, genotype, and sex at **a** P21 **b** P45 **c** P60 and **d** P80. Red represents depleted bacteria and blue represents sustained bacteria; color intensity reflects association strength. Significance was inferred by MaAsLin2 analysis using Log10 taxon. Detailed sample sizes are reported in Supplementary file 1.

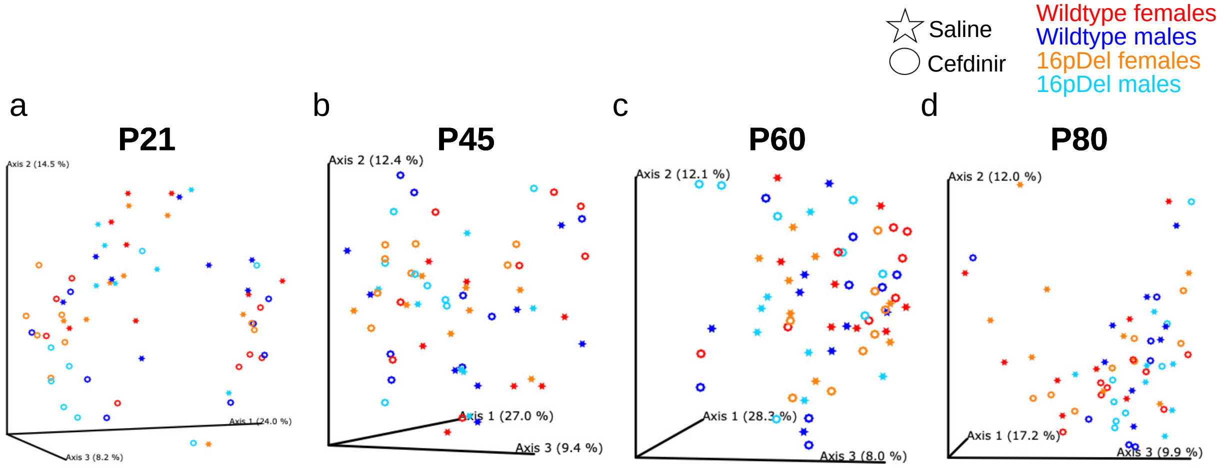


| | p value | q value |
|---|---|---|
| WT males | 0.162 | >0.05 |
| WT females | 0.069 | >0.05 |
| 16pDel males | 0.033 | >0.05 |
| 16pDel females | 0.098 | >0.05 |

| | p value | q value |
|---|---|---|
| WT males | 0.75 | >0.05 |
| WT females | 0.15 | >0.05 |
| 16pDel males | 0.437 | >0.05 |
| 16pDel females | 0.176 | >0.05 |

| | p value | q value |
|---|---|---|
| WT males | 0.652 | >0.05 |
| WT females | 0.407 | >0.05 |
| 16pDel males | 0.047 | >0.05 |
| 16pDel females | 0.623 | >0.05 |

| | p value | q value |
|---|---|---|
| WT males | 0.224 | >0.05 |
| WT females | 0.142 | >0.05 |
| 16pDel males | 0.461 | >0.05 |
| 16pDel females | 0.131 | >0.05 |

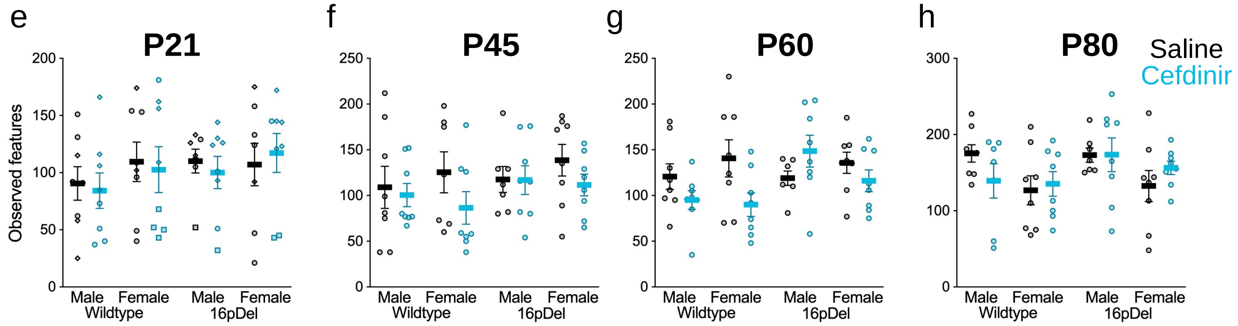


**Supplemental figure 4 (related to figure 1). Cefdinir-induced diversity reductions subside during the postweaning period.**

**a-d** PCoA plots showing Jaccard distances for individual fecal microbiota samples at **a** P21 **b** P45 **c** P60 and **d** P80. **e-h** Fecal $\alpha$-diversity, represented by the number of observed features, assessed across groups at each timepoint. Comparisons for **a-d** were performed using PERMANOVA pairwise tests with 999 permutations, with *p* and *q* values provided in the tables. Comparisons for **e-h** were made using multiple Mann Whitney tests with the two-stage step-up method of Benjamini, Krieger, and Yekutieli false discovery rate (FDR $q$ =1%). Data are presented as mean ± SEM. Detailed sample sizes are reported in Supplementary file 1.

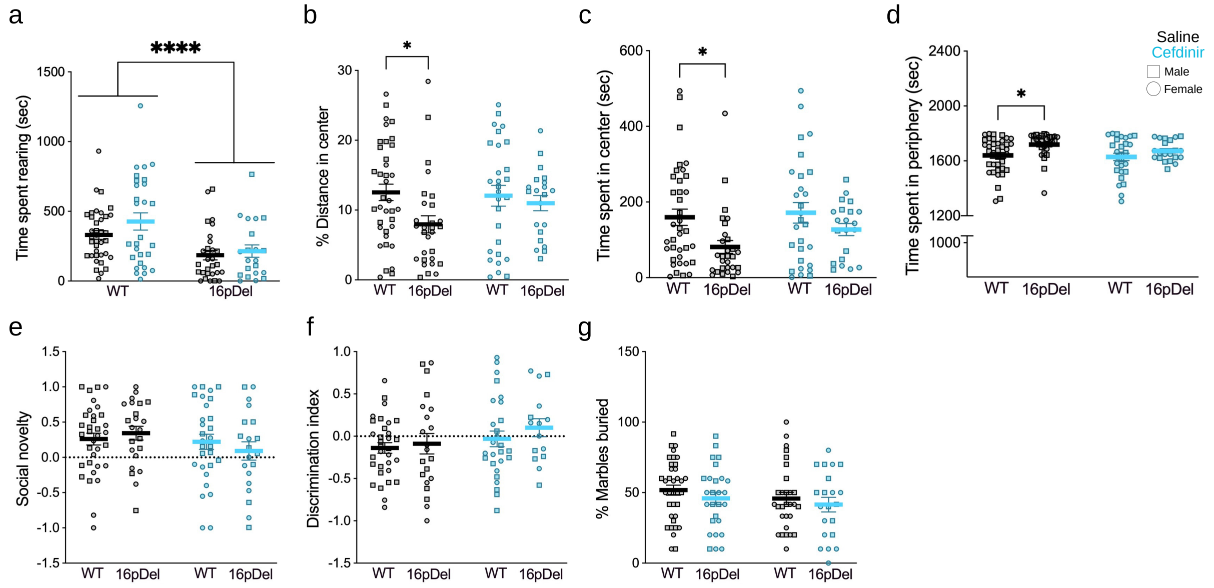


**Supplemental figure 5 (related to figure 2). Extended behavioral profiles across the lifespan.**

**a** 16pDel mice exhibited decreased exploratory (rearing) behavior compared to wildtype mice, independent of treatment F (1, 111) = 16.60, ****$p$<0.001. **b-d** At baseline, 16pDel mice displayed more anxiety-like behavior than wildtype littermates, evidenced by **b** decreased percent distance traveled in the center arena (F (1, 111) = 4.65, $p$=0.033, Tukey's adjusted $p$=0.041) **c** reduced time spent in the center (F (1, 111) = 7.67, P=0.007, Tukey's adjusted p=0.038) and **d** increased time spent in periphery (F (1, 111) = 7.67, p=0.007, Tukey's adjusted p=0.038). **e** Social novelty was comparable between saline- and cefdinir-exposed mice. The social novelty index was defined as the difference in time spent with the novel versus familiar mouse (from the social preference trial), divided by the sum of time spent with both. Positive values indicate preference for the novel mouse, whereas negative values indicate preference for the familiar mouse. **f** Short-term (90-min) recognition memory was unaffected by early-life cefdinir exposure. The discrimination index was calculated as the difference in time spent exploring the novel versus familiar object (from the familiarization trial) divided by the sum spent with both. Positive scores indicate recognition of novelty, while negative scores suggest a lack of object discrimination. **g** The percentage of marbles buried in the marble burying task did not differ between groups. Comparisons for **a-g** were made using a two-way ANOVA with Tukey's multiple comparisons test. All data are presented as mean ± SEM. Detailed sample sizes are reported in Supplementary file 1.

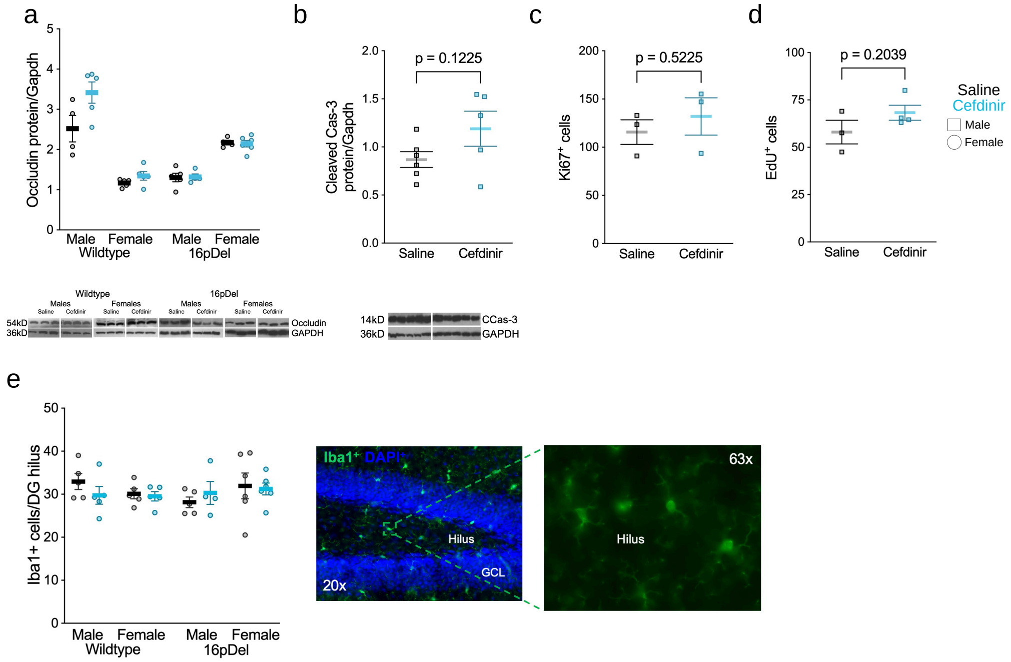


**Supplemental figure 6 (related to figure 3). Proliferative changes in the hippocampus are not due to blood brain barrier permeability or apoptosis.**

**a-b** Western blot analysis of **a** occludin and **b** cleaved caspase-3 protein levels, normalized to GAPDH, from individual 35µg whole brain lysates of P11 mice. Representative bands from a subset of mice in each group are provided. **c-d** Quantification of proliferative cells in the hilus and subgranular zone at P6, assessed by immunostaining for **c** Ki67$^{+}$ and **d** EdU$^{+}$ cells in saline- and cefdinir-exposed 16pDel males. **e** Quantification of Iba1$^{+}$ microglial cells in the hilus at P21. Comparisons for **a, e** were assessed using multiple Mann Whitney tests with the two-stage step-up method of Benjamini, Krieger, and Yekutieli false discovery rate (FDR *q* =1%). Comparisons for **b-d** were assessed using a Student’s t-test. All data are presented as mean ± SEM. Detailed sample sizes are reported in Supplementary file 1.

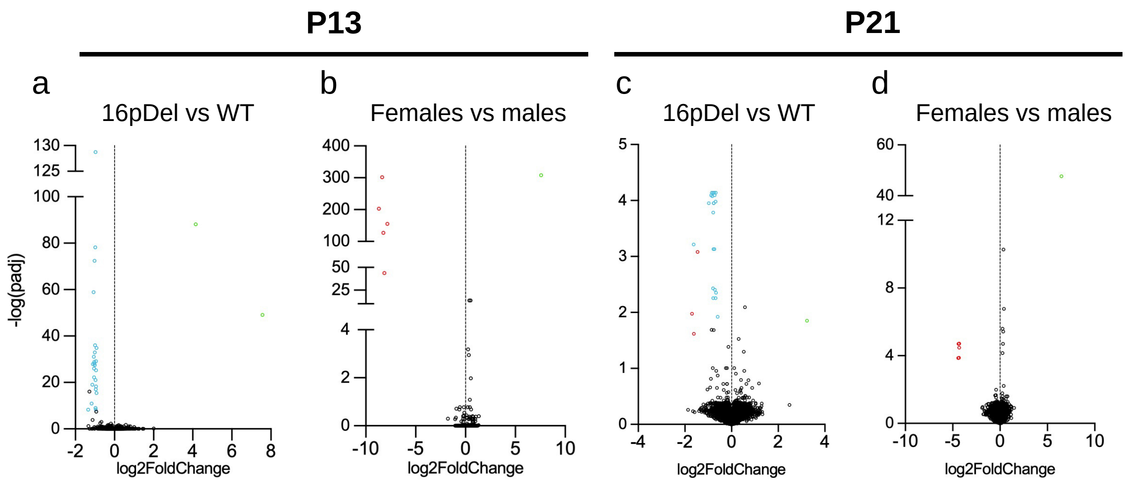


**Supplemental figure 7 (related to figure 4). Confirmation of expected differentially expressed genes across sex and genotype.**

**a-d** Volcano plots showing differentially expressed genes (DEGs) by **a,c** genotype and **b,d** sex at P13 and P21, respectively. DEGs were identified using the DeSeq2 package in R, with the Wald test applied to compute p-values and log2fold changes. Significance was defined as adjusted $p<0.05$ and log2fold <-1 or >1. Upregulated DEGs are depicted in green, downregulated DEGs in red, and nonsignificant genes (padj>0.05) in black. Genes within the 16p11.2 interval are depicted in cyan. Detailed sample sizes are reported in Supplementary file 1.
